# Recalling a single object: going beyond the capacity debate

**DOI:** 10.1101/2020.02.17.951921

**Authors:** Rakesh Sengupta, Christelle M. Lewis, Raju S. Bapi

## Abstract

Working memory is now established as a limited capacity system. The debate regarding working memory has been largely between slots and resource based models. The resource model suggests that as the number of items increases, precision of recall decreases because neural resources are dynamically allocate to all the objects needed for task. Slot model on the other hand implies that an item is stored either with the highest precision or not at all. If both these models stand true then quality of memory performance would be near perfect for a single object. However, that may not be the case. In the current work, we investigated recall accuracy for feature(s) of a single object in three successive experiments. In all three experiments, the memory array consisted of a single colored oriented short line presented a short distance away from the center of the display for 1 sec. We probed recall of features from the set of color, location, orientation, and size. In experiment 1 number of recall question varied between 1 – 4 with the order randomized in each trial. In experiment 2 we chose to probe only two feature recall questions, whereas only one recall question was probed for the third experiment. In experiment 3 we varied the delay before the recall probe between 1 and 2 s. The recall response for each feature was mapped on to a continuous variable. Subjects used a color wheel to respond to color, on-screen mouse click to indicate center of the line location, click away from the center to indicate size, and a mouse click to the periphery of centered circle to indicate orientation with the slant of the resulting radial line. We calculated z-scores of errors for each feature for every subject separately. In experiment 1 that for color, location, and size the errors increase significantly with the position of the questions asked. In experiment 2, the errors increased significantly between questions for color and location (but not for orientation and size). In experiment 3, we did not see any significant increase in error with recall probe delay. Overall run-time for each trial was within 10 secs, well within the limits of operation of working memory. This drop in performance poses questions for memory mechanisms proposed by slot and resource models as both would predict near-perfect recall within the time-period for the trials.

## Introduction

Following the conceptualization and seminal contribution by Baddeley and Miller, working memory has entered the lexicon of cognitive psychology as one of the key executive functions of the brain. We invoke the concept of working memory to understand how brain manages to perform tasks that are not resolved by the current percept alone and requires manipulation and storage of information temporarily. The major theoretical frameworks currently available for working memory in general are influenced by largely the following categories (note that not all the following are mutually incompatible)

- Multi-component model of working memory where a central executive directs a phonological loop, visuo-spatial sketchpad, and episodic buffer for maintenance and manipulation of temporary task related information(Baddeley, 2003, 2012).
- Working memory as a maintenance of representations from long term memory that are activated within the focus of attention (Cowan, 2001).
- Working memory as a limited capacity maintenance system for discrete fixed-resolution items needed for task completion (Luck and Vogel, 1997; Zhang and Luck, 2008).
- Working memory as a limited resource distributed dynamically and flexibly over all the items that need to be maintained in memory (Ma et al., 2014).

Last couple of decades has seen a narrowing of the focus of working memory research towards psychological and neurophysiological studies of capacity (for an extensive review see Cowan (2012) and Luck and Vogel (2013)).The research on the capacity of visual working memory (VWM) has largely focused on a slot based approach (e.g. fixed slots (Luck and Vogel, 1997)), or approaches with a non-specific neural architecture (e.g. dynamic resource allocation with varied precision, among others(Ma et al., 2014)). Interestingly, the debate has predominantly been within these paradigms, and has progressed for the most part without a firm proposal regarding the actual content of VWM or the actual mechanisms of recall. Thus, researchers probe capacity limit of VWM with paradigms that involve multiple objects presented one at a time in sequence (serial recall) or multiple objects presented simultaneously in a brief memory array that can be probed in a test array (e.g., change detection paradigms) or with location or feature probes.

However, we propose that such paradigms for VWM leave some crucial mechanistic aspects in the dark.

1. What is the actual content of VWM?
2. How does the process of recall affect the content of VWM?

We decided to probe these questions with a series of experiments involving the recall of 1 or many (maximum 4) features (color, location, size, and orientation) of a single object. According to both slot and resource models, since there is one single object to be held in memory, recall should be sufficiently noiseless (at least for the duration of maximum 8-10 secs till all the questions are asked) in absence of any other item to overload the capacity of slots or to stretch the available resources.

The structure of all three experiments presented in the paper followed a simple paradigm involving recall of features one after one from a memory array consisting of a single object - a short, colored, oriented line presented at a certain distance from the center of the screen. No. of feature recall questions varied from experiment to experiment. The features probed for recall were color, location (of the center of the line), orientation, and size. In the next sections we will elaborate on the methods and the results for each experiment.

## Experiment 1

### Participants

Twenty healthy undergraduate students and staff (11 male, 9 female; all right handed with an average age of 22 years) with normal to corrected-to-normal vision, normal color perception, were recruited from personal of SR Engineering College. Informed consent was obtained from all the participants before commencing the experiment.

### Apparatus

Participants performed the experiments in a dark room. The position of each participant’s head was placed 60 cm away from an LCD The participants were given instructions and then seated in front of the experimental system in a dark room. The participant’s head was stabilized 60cm away from the an LCD screen (34.5cm*19.4cm, resolution: 1366*768, refresh rate: 60.00Hz). Experiment was designed in Psychtoolbox and was maintained by Octave 4.4.1.

### Stimuli

Short colored rectangular lines (Width: 5 pixels) of varying length and orientation were used as the stimulus. From trial to trial, the stimulus appeared in random locations across the screen. Only one rectangular bar was shown per trial. The size of the stimuli was varied between 1 to 3 degrees of visual angle, The location was varied from 0 to 4 degrees visual angle from the center, and the orientation was varied across 360 degrees. Their colors were randomly chosen from an equi-luminant (L=50) CIELAb color space. The conversion from the lab space is based on Bruce Lindbloom’s excellent website about color spaces.

### Design

The experiment began when the subjects were ready. They were presented with a red central fixation cross for 500ms on a white screen that signaled the start of every trial. In each trial, a single oriented colored line of varying size appeared anywhere across the screen and stayed so for 1000ms. Across each trial the features color, location, orientation and size of the stimuli varied. After a 1000ms retention period, the subjects were asked a few questions about the features of the stimuli. They had to recall all or some of the features of the line. A maximum of 4 questions were asked across 120 trials. They judged what the color of the stimuli was with a mouse click from a color wheel consisting of 360 graduation of the primary color on the circumference of the circle. The colors were chosen from the CIELab Color space (*L* = 50). The distance between the color shown and the color selected was taken as the error for the color response. To determine the orientation the line, the subjects were asked to select a point on a black circle such that the line from the center of the circle to the selected point on the circle was parallel to the stimuli. The difference between the angle of the stimuli and the angle of the response line was considered as the error for the orientation response. For locating where the stimuli appeared, a blank screen was presented and the subjects were required to click anywhere on the screen where the midpoint of the line supposedly was. The difference between the stimulus’ midpoint and the selected point was taken as the error in determining location. To deduce the length of the line, a black cross appeared at the center of the screen as an inference for the starting point of the line. The subjects were asked to click at a point away from the central cross such that the distance between the two points would be equal to the size of the stimuli shown in the trial. The difference between the the length of the stimuli and the recalled size of the line was considered as the error in size. The questions were asked in a random order one after the other and remained on screen until a response was recorded from the mouse click. A blank white screen was presented for 2000 ms after the answers were given. The total of 120 trials took 20 minutes to complete. No breaks were given in between the experiment. The design is represented in Fig. 1.

**Figure 1.**
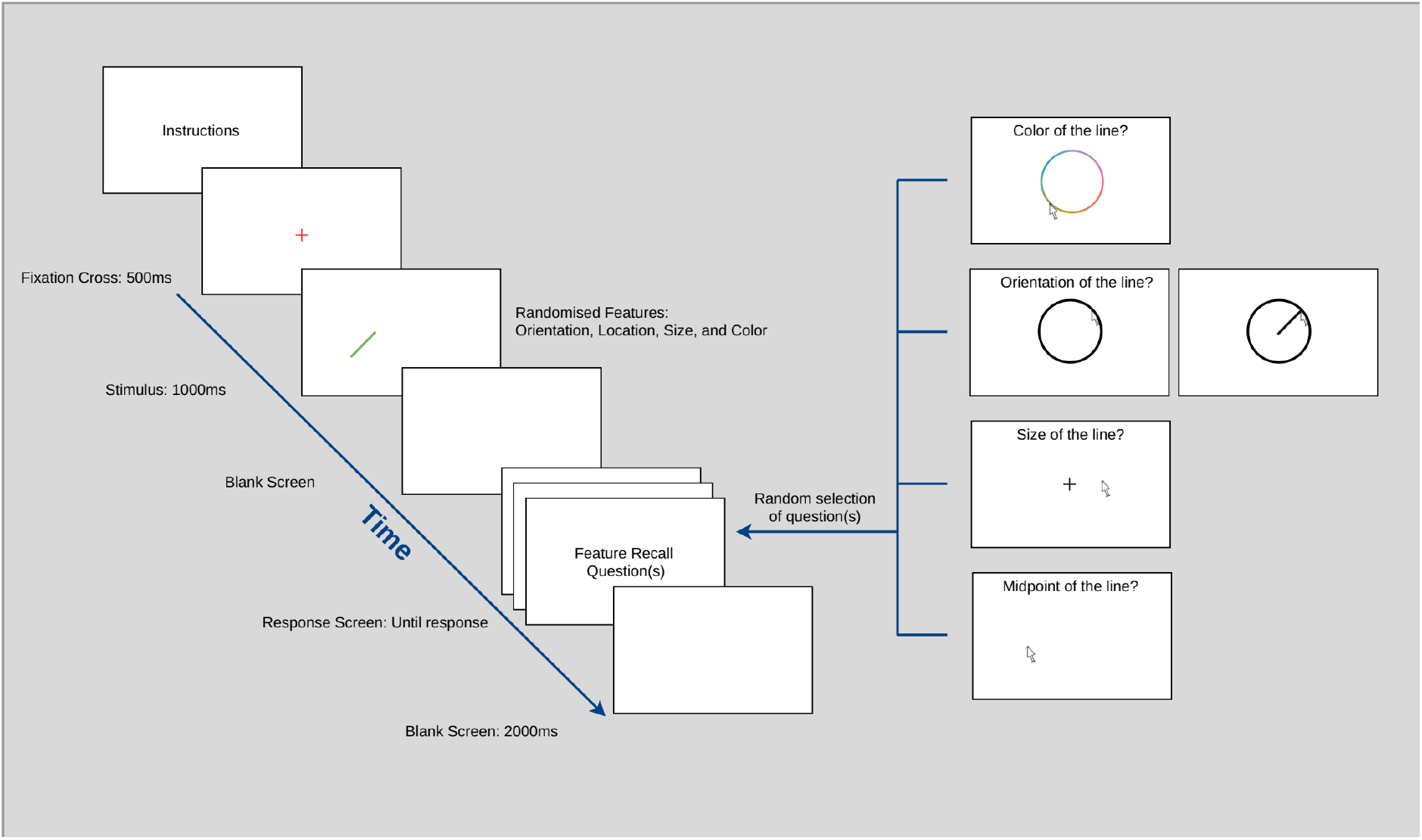
Design of the experiments. Subjects began every trial with a fixation cross presented for 500 ms. Then a short, colored, oriented line was presented at a variable distance at the center of the screen for a duration of 1s. A blank screen was shown during retention period. The retention period before the recall question was ~ 1s before the first question for Experiments 1 & 2. The recall questions asked subjects to specify color, location, orientation, and size of the remembered line. In Experiment 1 the number of recall question varied between 1 and 4, whereas in Experiment 2 the number of recall questions was fixed to 2. In Experiment 3 only one feature was probed for recall in each trial and the retention interval was varied between 1s and 2s (a ±.1s jitter was applied to both the intervals). The number of trials for Experiments 1, 2, and 3 were 120, 160, and 160 respectively. The order of feature recall questions were randomized in each trial.

### Results

The raw error values for each feature corresponding to the serial position of the corresponding question are given in Table A1. The standard errors are reported in parenthesis after the means. Color errors are reported as angular distance from the expected color on the color wheel. Location and size errors are reported in degrees of visual angles. Orientation errors are reported in angular distance as well. For size errors we calculated absolute values.

Before the analysis, we converted the errors (for each subject and each feature separately) into their corresponding z-scores by subtracting the mean error for the particular feature and dividing by their standard deviation.

The results shown in Fig. 2 shows that for color, location, and size the errors increase if the corresponding feature query is asked later in the response sequence, but not for orientation. We conducted a two-way within subject repeated measures ANOVA for the *z*-scores of the errors. We found main effect of position of feature question (*F*(3,228) = 48.79, *p* < 0.001, *η*^2^ = 0.32, 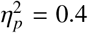) as well as a significant interaction between features and position of question (*F*(9,228) = 8.14, *p* < 0.001, *η*^2^ = 0.16, 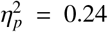). The post-hoc analysis was carried out with a series of 1-way ANOVA tests for each feature (for instance between error in color recall between positions 1 and 2, 2 and 3, 3 and 4 etc and). The results are depicted in Fig. 3. For details of post-hoc statistics see Table B1.

**Figure 2.**
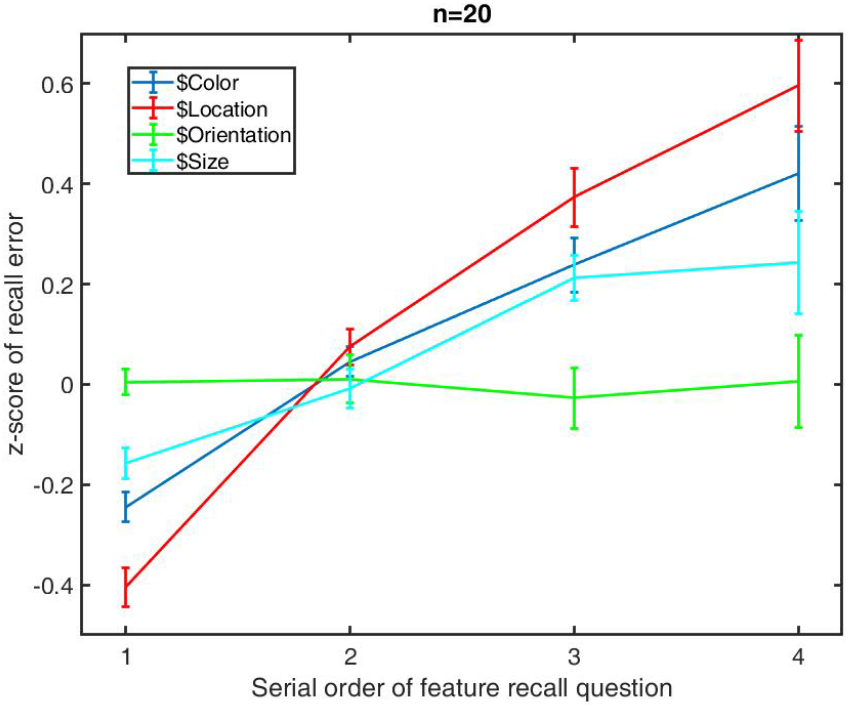
Average *z*-scores across the participants for each feature question in the order of the serial position of the question asked in Experiment 1.

**Figure 3.**
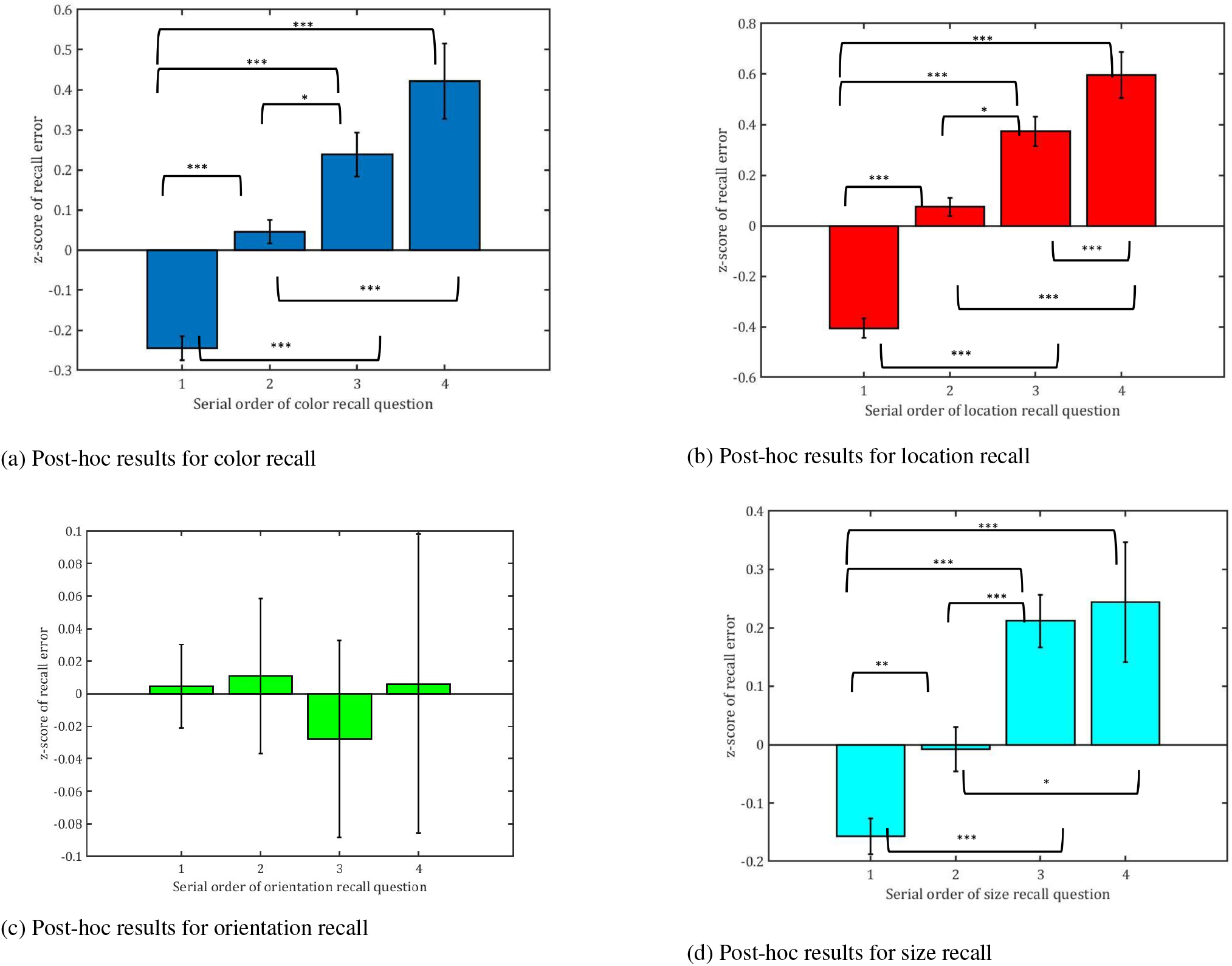
Post-hoc results for each feature probed for recall. We can that color, location, and size show increasing recall errors with the serial position in which the recall question was probed for.

In the current experiment we tested the recall of single object (colored oriented line) of a particular size and presented within a short distance from the center of the screen. Number of feature questions to be asked was varied on trial-to-trial basis and so was the order of questions (regarding color, location, orientation, or size of the object). The total trial length for even the four questions did not exceed 10-15 seconds, well within the limits of operation of working memory. The second question was asked within 3-4 s of the stimulus. If one subscribes to most standard ideas of VWM (slots, resources, or other capacity based ideas), there should not have been an increase in error with the position of feature question. However, we found a significant increase in error.

## Experiment 2

One explanation for the drop in accuracy with increasing number of questions can be due to cognitive load and uncertainty regarding the potential number of feature questions. Thus in the second experiment we chose to limit the number of questions to just 2 per trial. Every other experimental parameters for Experiment 2 remained the same as that of Experiment 1. The no. of trials was changed to 160 per participants. 20 subjects (12 male and 8 female; avaerage age 21) chosen from SR Engineering College, Warangal participated in the experiment.

### Results

The mean recall error values for Experiment 2 are reported in Table A2 and graphical results are depicted in Fig. 4. Results of within-subject repeated measures ANOVA (conducted after conversion of errors to their corresponding z-score - like in Experiment 1) shows a strong main effect for position of feature question (*F*(1,76) = 110.56, *p* < 0.001, *η*^2^ = 0.42, 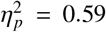) as well as a significant interaction between features and position of recall questions (*F*(3,76) = 26.24, *p* < 0.001, *η*^2^ = 0.3, 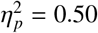).

**Figure 4.**
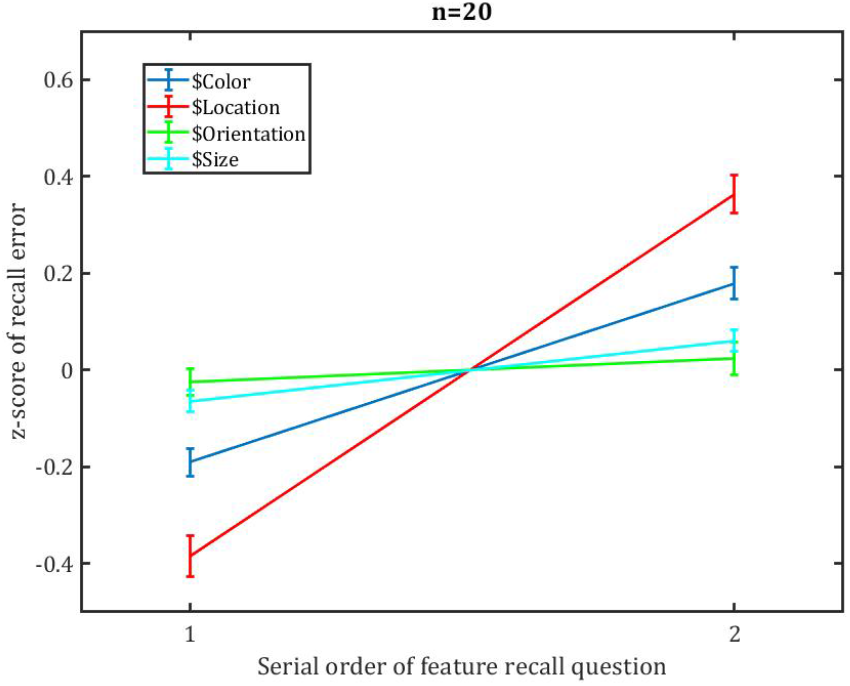
Average z-scores across the participants for each feature question in the order of the serial position of the question asked in Experiment 2.

We conducted post-hoc test for each feature separately here the mean errors for first position in recall order were compared with errors for second position in the recall order using simple 1-way ANOVA. The results replicated the findings from Experiment 1. Color (*F*(1, 38) = 75.57, *p* < 0.001), location (*F*(1,38) = 167.00, *p* < 0.001), and size (*F*(1,38) = 16.47, *p* < 0.001) show increase in recall error between serial position 1 and 2, as opposed to orientation (*F*(1,38) = 1.28, *p* = 0.26).

In this experiment we see the same pattern of results repeated as depicted in Fig. 2 for Experiment 1. Thus we can rule out the possibility of extraneous cognitive load from uncertainty regarding number of questions, or the number of questions themselves leading to the increase in error for features like color, location, and size.

## Experiment 3

A plausible explanation may be provided for the results so far in the delay in the questions’ appearances themselves. It might be possible that the second question was asked too late (although the actual delay might be as little as 2-3 sec). To explore this possibility, we conducted a third experiment where we asked only one out the four possible feature recall question after the retention interval in each trial, but we manipulated the retention interval. Total of 15 subjects (10 male and 5 female; average age of 23) participated in this experiment.

### Design

In this experiment the stimuli were the same as in previous 2 experiments. The novel aspect in this experiment was the manipulation of the retention interval before the single feature recall question per trial. The retention interval was randomly chosen from two interval windows centered around 1 and 2 sec respectively. A jitter of ±0.1 or ±0.05 s was added randomly in each trial. No. of participants was 15 and total number of trials was 160.

### Results

The results for Experiment 3 are depicted in Table A3 and Fig. 5. We can see that with increased retention interval there is no significant increase in error scores. After converting feature specific error scores to the corresponding z-scores, we found no significant effect of retention interval (*F*(1,56) = 1.15, *p* = 0.23, *η*^2^ = 0.02, 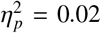). We also did not find any significant interaction term (*F*(3,56) = 0.6, *p* = 0.61, *η*^2^ = 0.03, 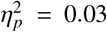). As there was no main effect, post-hoc tests were not performed.

**Figure 5.**
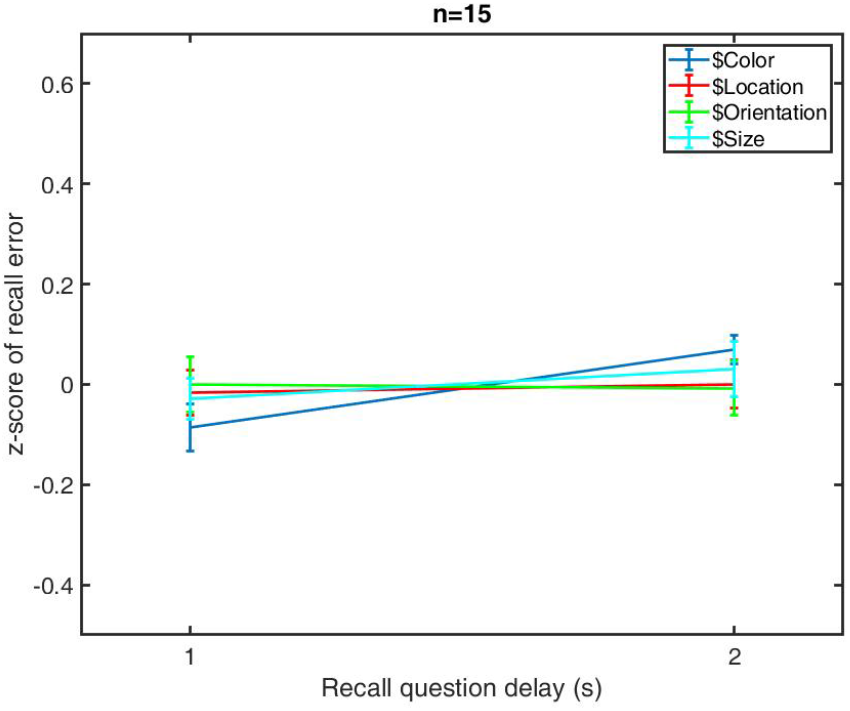
Average z-scores across the participants for each feature question in the order of the serial position of the question asked in Experiment 3.

## Discussion

In the series of experiments presented previously we investigated the recall of single item. Recall was tested with a series of feature recall questions in Experiments 1 & 2. In both we saw (see Fig. 2 and 4) that recall of features like color, location, and size suffer a recall penalty if that feature is asked for later in the sequence of questions. We also confirmed through the manipulation of retention interval in Experiment 3 (see Fig. 5) that this increase in the error of recall is not an artifact of temporal distance of the recall question from the stimulus.

The conventional capacity based explanations lose their value in this experiment as we have only one object in the memory array that stays on screen for 1 s. Moreover, we tested for recall rather than recognition, thus the recall error was mapped to continuous variables. The standard models say that working memory is supposed to keep task-relevant information until the task is completed. Here the subjects did not know how many questions they needed to answer beforehand. Thus before task completion, there should have been no need for memory reset allowing for increased error. The contents of VWM should also not be subject to forgetting similar to Long-Term memory either.

If slot model can be accepted, then either one remembers the entire object or forgets the entire object. We should not see the gradual trend of feature recall errors with increasing number of questions as we see in Fig. 2. However, one can somewhat argue in the favor of resource model, that maybe if the subjects deplete their neural resources during the first question, they may show poorer performance at the subsequent questions. In order to probe the possibility we checked for pairwise feature recall errors in Experiment 2. We chose experiment 2 because in this experiment we probed only 2 questions per trial. For this purpose we isolated three kinds of trials - a) trials where only color and location were probed, b) trials where only color and size were probed, c) trials where only location and size were probed. We excluded orientation trials from this analysis as orientation recall error did not show variation between conditions. For each trial type, we further separated into ‘color first’ and ‘location first’ (for trial types a), ‘color first’ and ‘size first’ (for trial type b) trials and so on. For trial type a we checked for correlation of recall error between color and location trials for both ‘color first’ and ‘location first’. We performed similar analysis for the other trial types as well. According to resource models we should expect negative correlations for all of them. However, we can see from Fig. 6,7,8 that such is not the case. In fact quite a few of the subjects show a positive correlation between them.

**Figure 6.**
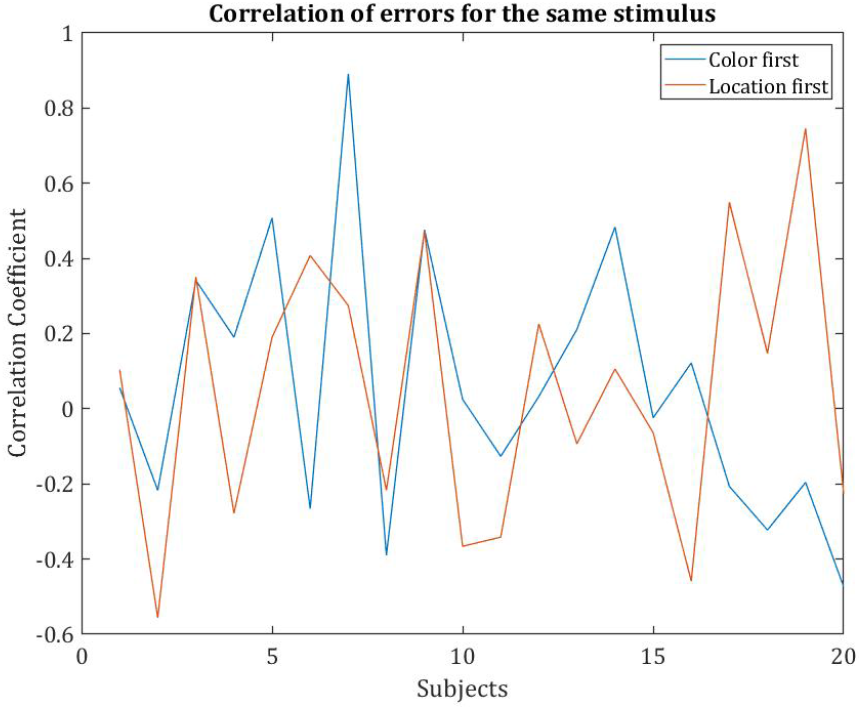
Subject-wise correlation between color and location recall errors in color-location trials in experiment 2.

A possible explanation for our results can be an effect similar to inhibition-of-return that kicks in when a subject answers a question regarding the content of VWM. It is similar to the real world phenomena where a person being introduced to another for the first time immediately forgets the name of other as soon as the formalities of introduction are concluded. In our future work we intend to investigate the origins of such a possible mechanism in greater depth. Moreover, we can see that there is a room for further exploration into individual differences of recall errors, as Fig. 6,7,8 suggest that some subjects might be equally good or bad at recall, and there might also be different strategies employed by different participants.

**Figure 7.**
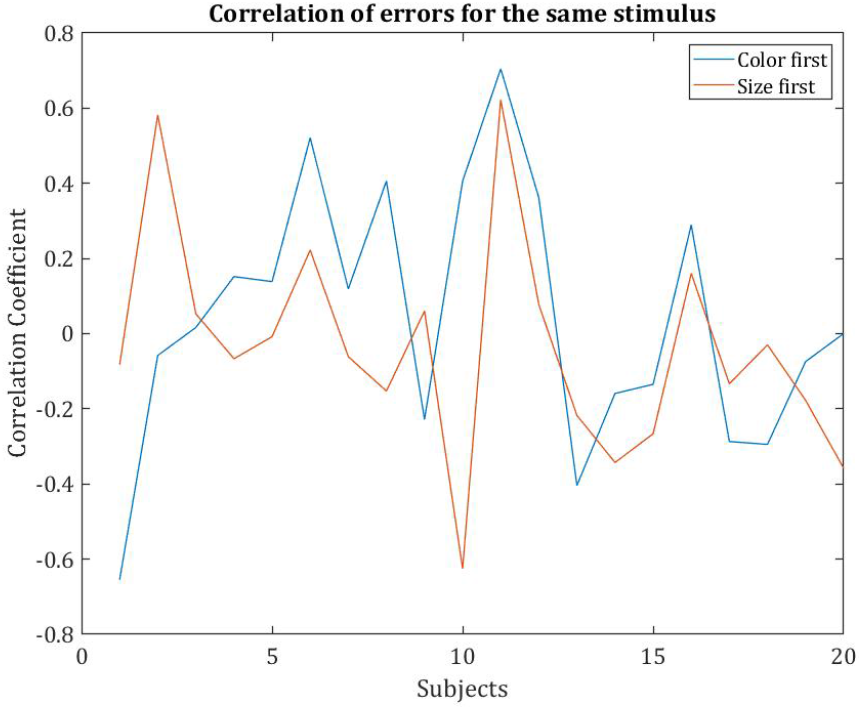
Subject-wise correlation between color and size recall errors in color-size trials in experiment 2.

**Figure 8.**
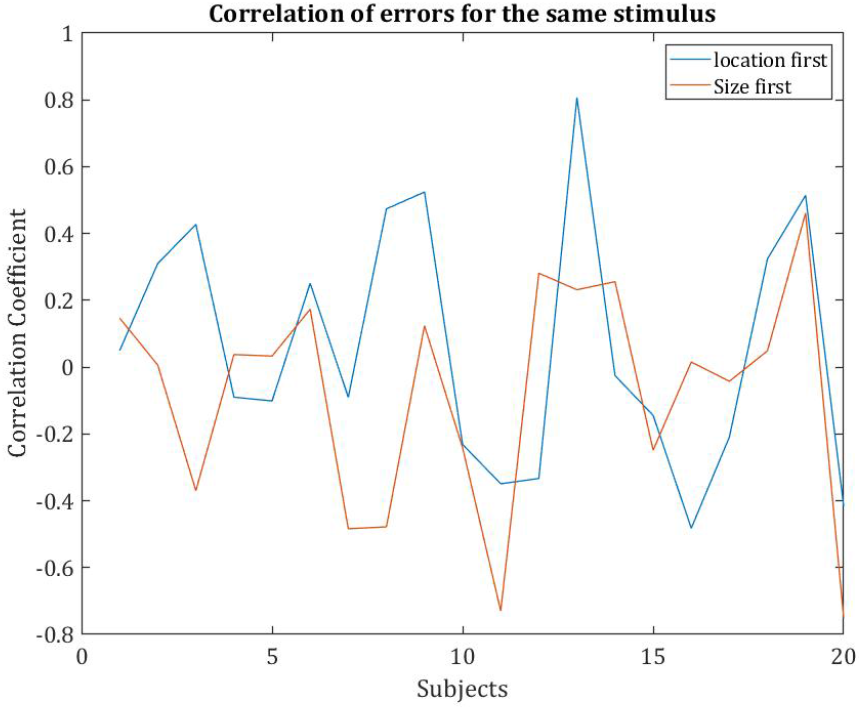
Subject-wise correlation between size and location recall errors in size-location trials in experiment 2.

A crucial aspect of the results show differentiation between recall of features like orientation from others. Not all features suffer the recall error penalty. This might point to the difference between features that are encoded in different stages of executive function of VWM.

This brings to question the passive container metaphor for working memory that all current theories subscribe to. We should think of a more spontaneous and active executive processes of working memory.

## Appendix A Raw results

**Table A1.** Raw recall errors for each feature in Experiment 1. The errors in color recall are reported in angular distance in color wheel (in degrees). Location and size errors are reported in degrees of visual angle. Orientation errors are reported in degrees of angular distance.

|  |  | Serial position of feature question |  |  |  |
| --- | --- | --- | --- | --- | --- |
|  |  | 1 | 2 | 3 | 4 |
| Feature recall error | Color (in °) | 17.32 (± 0.91) | 23.79 (± 1.17) | 29.06 (± 2.50) | 33.10 (± 3.11) |
|  | Location (in °) | 0.72 (± 0.07) | 1.11 (± 0.12) | 1.36 (± 0.15) | 1.49 (± 0.15) |
|  | Orientation (in °) | 58.75 (± 3.14) | 58.72 (± 2.79) | 57.59 (± 3.86) | 59.01 (± 3.82) |
|  | Size (in °) | 0.43 (± 0.06) | 0.48 (± 0.07) | 0.56 (± 0.07) | 0.56 (± 0.08) |

**Table A2.**
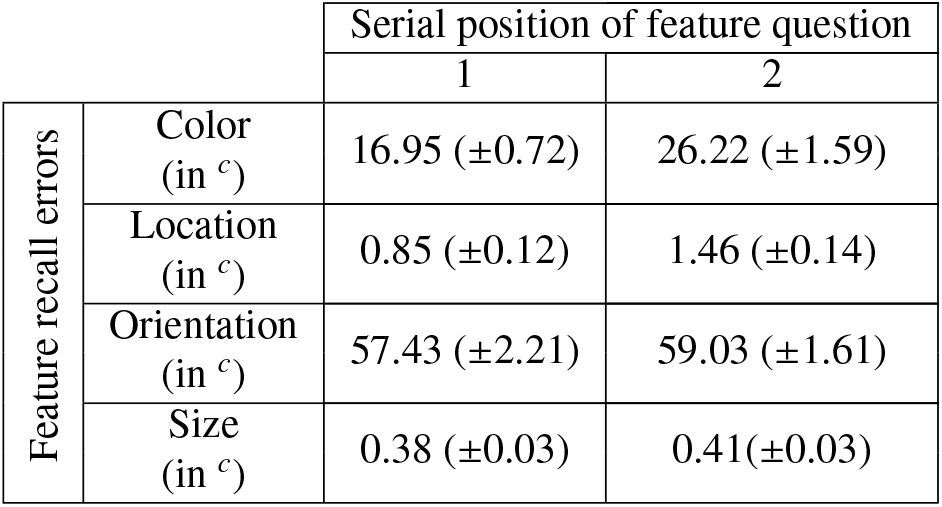
Raw recall errors for each feature in Experiment 2.

**Table A3.**
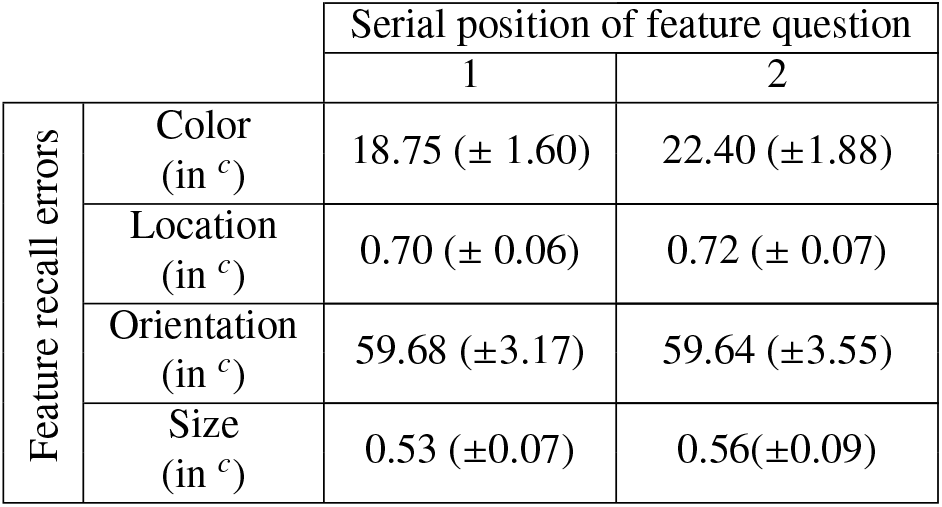
Raw recall errors for each feature in Experiment 3.

## Appendix B Post-hoc results

**Table B1.** Post-hoc results for Experiment 1

|  | 1-2 | 1-3 | 1-4 | 2-3 | 2-4 | 3-4 |
| --- | --- | --- | --- | --- | --- | --- |
| Color | $F(1, 38) = 48.23$,<br>$p < 0.001$ | $F(1, 38) = 37.3$,<br>$p < 0.001$ | $F(1, 38) = 23.83$,<br>$p < 0.001$ | $F(1, 38) = 9.54$,<br>$p < 0.05$ | $F(1, 38) = 8.29$,<br>$p < 0.01$ | $F(1, 38) = 2.82$,<br>$p > 0.1$ |
| Location | $F(1, 38) = 83.13$,<br>$p < 0.001$ | $F(1, 38) = 75.17$,<br>$p < 0.001$ | $F(1, 38) = 51.76$,<br>$p < 0.001$ | $F(1, 38) = 19.03$,<br>$p < 0.001$ | $F(1, 38) = 15.75$,<br>$p < 0.001$ | $F(1, 38) = 4.21$,<br>$p < 0.05$ |
| Orientation | $F(1, 38) = 0.014$<br>$p > 0.1$ | $F(1, 38) = 0.19$<br>$p > 0.1$ | $F(1, 38) = 0.08$<br>$p > 0.1$ | $F(1, 38) = 0.25$<br>$p > 0.1$ | $F(1, 38) = 0.09$<br>$p > 0.1$ | $F(1, 38) = 0.09$<br>$p > 0.1$ |
| Size | $F(1, 38) = 9.26$<br>$p < 0.01$ | $F(1, 38) = 23.24$<br>$p < 0.001$ | $F(1, 38) = 9.65$<br>$p < 0.001$ | $F(1, 38) = 13.74$<br>$p < 0.001$ | $F(1, 38) = 4.02$<br>$p < 0.05$ | $F(1, 38) = 0.08$<br>$p > 0.1$ |

